## Supplementary file 1 for "Endocrine disruptor-induced epimutagenesis *in vitro*: Insight into molecular mechanisms"

**Dose-specific differentially methylated sites (DMCs) (treated vs. control)**

| **DMCs** | **iPSC 1** μ**M** | **iPSC 50** μ**M** | **iPSC 100** μ**M** |
| --- | --- | --- | --- |
| Hypomethylated | 9,066 | 3,456 | 14,337 |
| Hypermethylated | 3,107 | 27,734 | 11,223 |
| Total | 12,173 | 31,190 | 25,560 |
