## Supplementary file 2 for "Endocrine disruptor-induced epimutagenesis *in vitro*: Insight into molecular mechanisms"

**Dose-specific differentially methylated regions (DMRs) (treated vs. control)**

| **DMRs** | **iPSC 1** μ**M** | **iPSC 50** μ**M** | **iPSC 100** μ**M** |
| --- | --- | --- | --- |
| Hypomethylated | 8 | 3 | 85 |
| Hypermethylated | 0 | 32 | 12 |
| Total | 8 | 35 | 97 |
