## Supplementary file 3 for "Endocrine disruptor-induced epimutagenesis *in vitro*: Insight into molecular mechanisms"

**Dose-specific differentially expressed genes (DEGs) (treated vs. control)**

| **DEGs** | **iPSC 1** μ**M** | **iPSC 50** μ**M** | **iPSC 100** μ**M** |
| --- | --- | --- | --- |
| Up regulated | 868 | 2,139 | 2,524 |
| Down regulated | 464 | 1,926 | 2,554 |
| Total | 1,332 | 4,065 | 5,078 |
