## Supplementary file 4 for "Endocrine disruptor-induced epimutagenesis *in vitro*: Insight into molecular mechanisms"

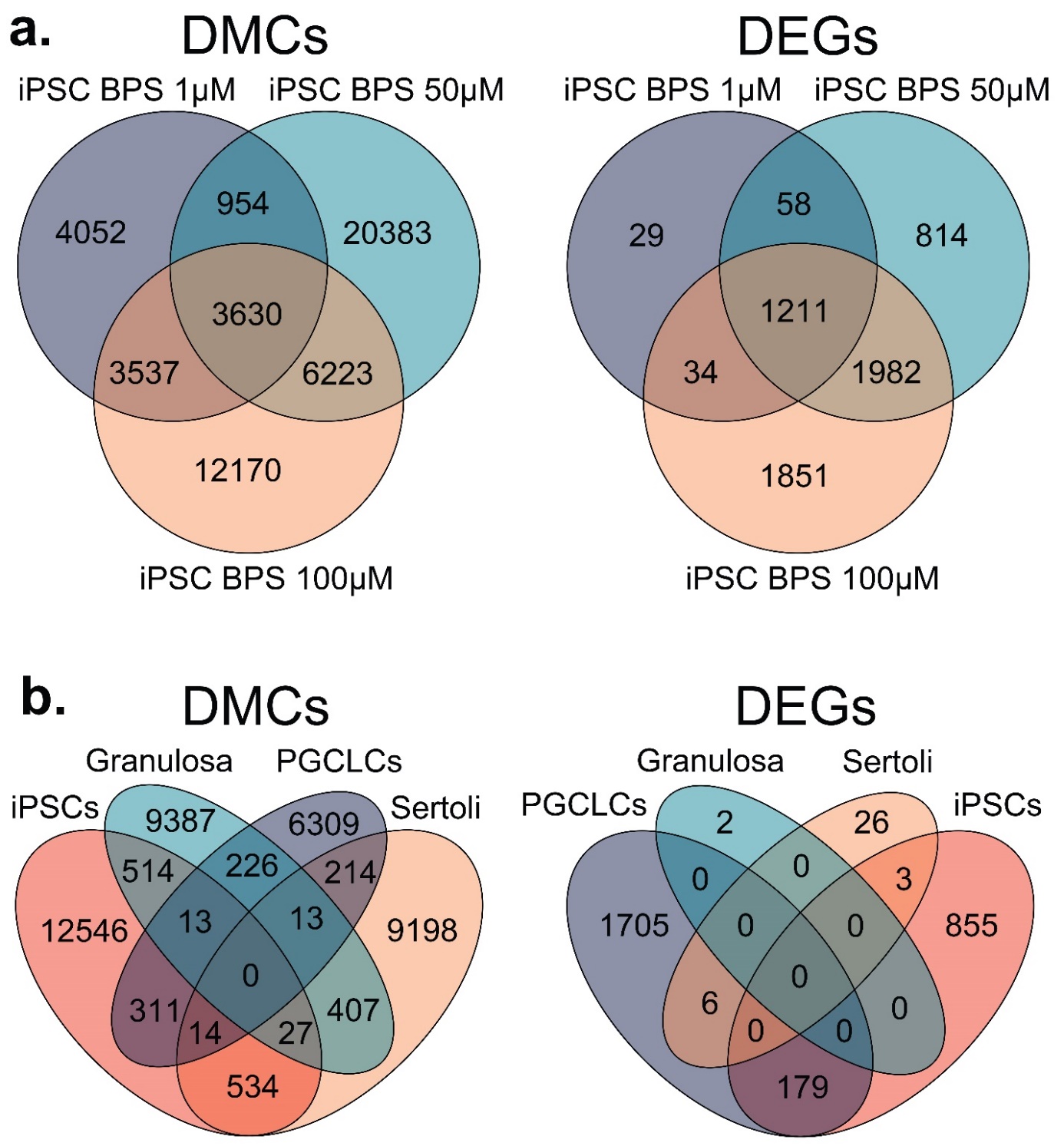

**Overlapping DMCs and DEGs found among dose-dependent and cell-type specific responses to BPS exposure.** Venn diagrams of overlapping DMCs/DEGs identified when comparing **a)** iPSCs exposed to increasing doses of BPS (1, 50, and 100 μM) or **b)** iPSCs, granulosa cells, Sertoli cells, and PGCLCs exposed to the established minimum dose of 1 μM of BPS. We detected an average of 51.25% overlap among DMCs and an average of 80.45% overlap among DEGs within each respective cell type across the different doses of BPS, but only an average of 11.05% among DMCs and 13.26% among DEGs between different cell types.
