## Supplementary file 5 for "Endocrine disruptor-induced epimutagenesis *in vitro*: Insight into molecular mechanisms"

**DEGs containing DMCs observed in iPSC exposed to increasing doses of BPS**

| **DEGs containing DMCs** | **iPSC 1** μ**M** | **iPSC 50** μ**M** | **iPSC 100** μ**M** |
| --- | --- | --- | --- |
| Promoter | 264 (19.82%) | 693 (17.04%) | 1,136 (22.37%) |
| Gene body | 436 (32.73%) | 1,541 (37.91%) | 1,934 (38.08%) |
