## Supplementary file 6 for "Endocrine disruptor-induced epimutagenesis *in vitro*: Insight into molecular mechanisms"

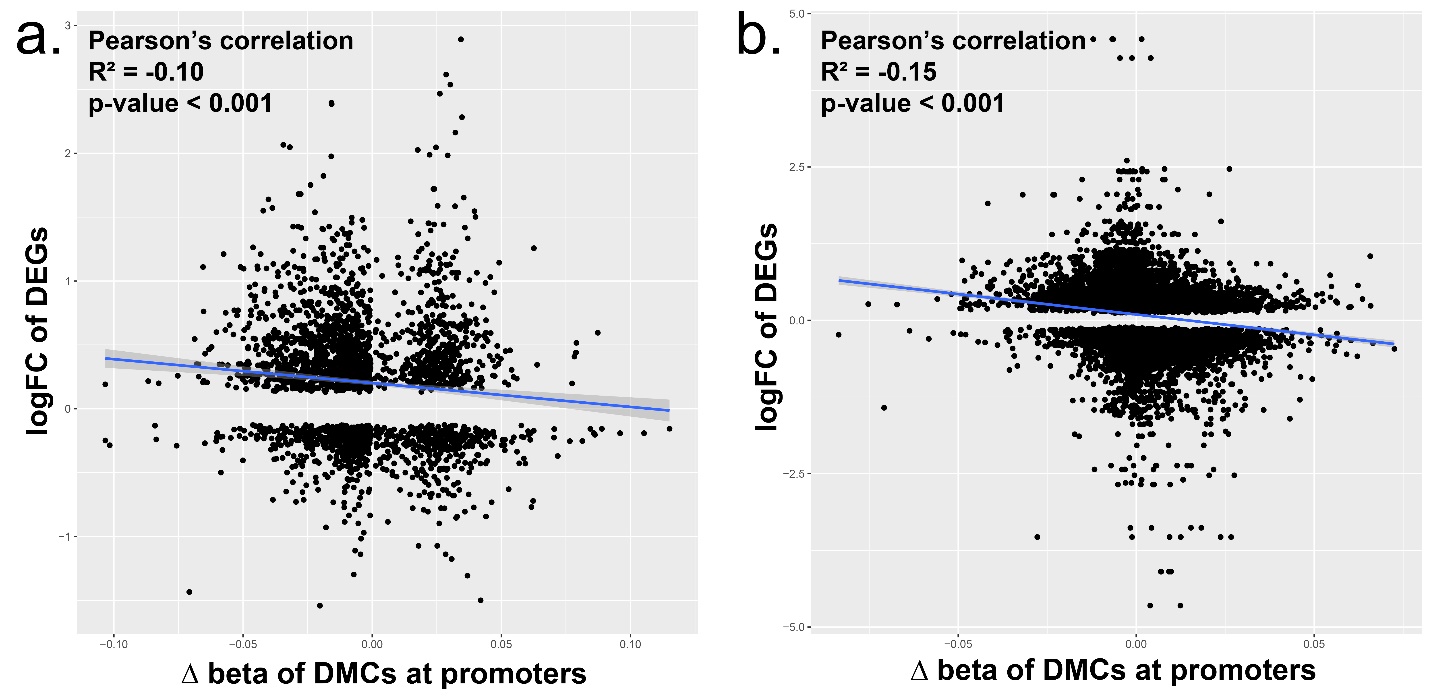

**Relationship between DMCs at promoters and DEGs.** We compared the correlation between the differential expression of genes and presence of DMCs in the promoter region of that gene in **a) )** iPSCs exposed to increasing doses of BPS (1, 50, and 100 μM) or **b)** Sertoli, Granulosa, iPSCs, and PGCLCs. We found that there was a significant negative correlation indicating in our data that promoters that had a loss of DNA methylation tended to also have higher upregulation of gene expression and vice versa when observing hypermethylation and downregulation of gene expression.
