## Supplementary file 7 for "Endocrine disruptor-induced epimutagenesis *in vitro*: Insight into molecular mechanisms"

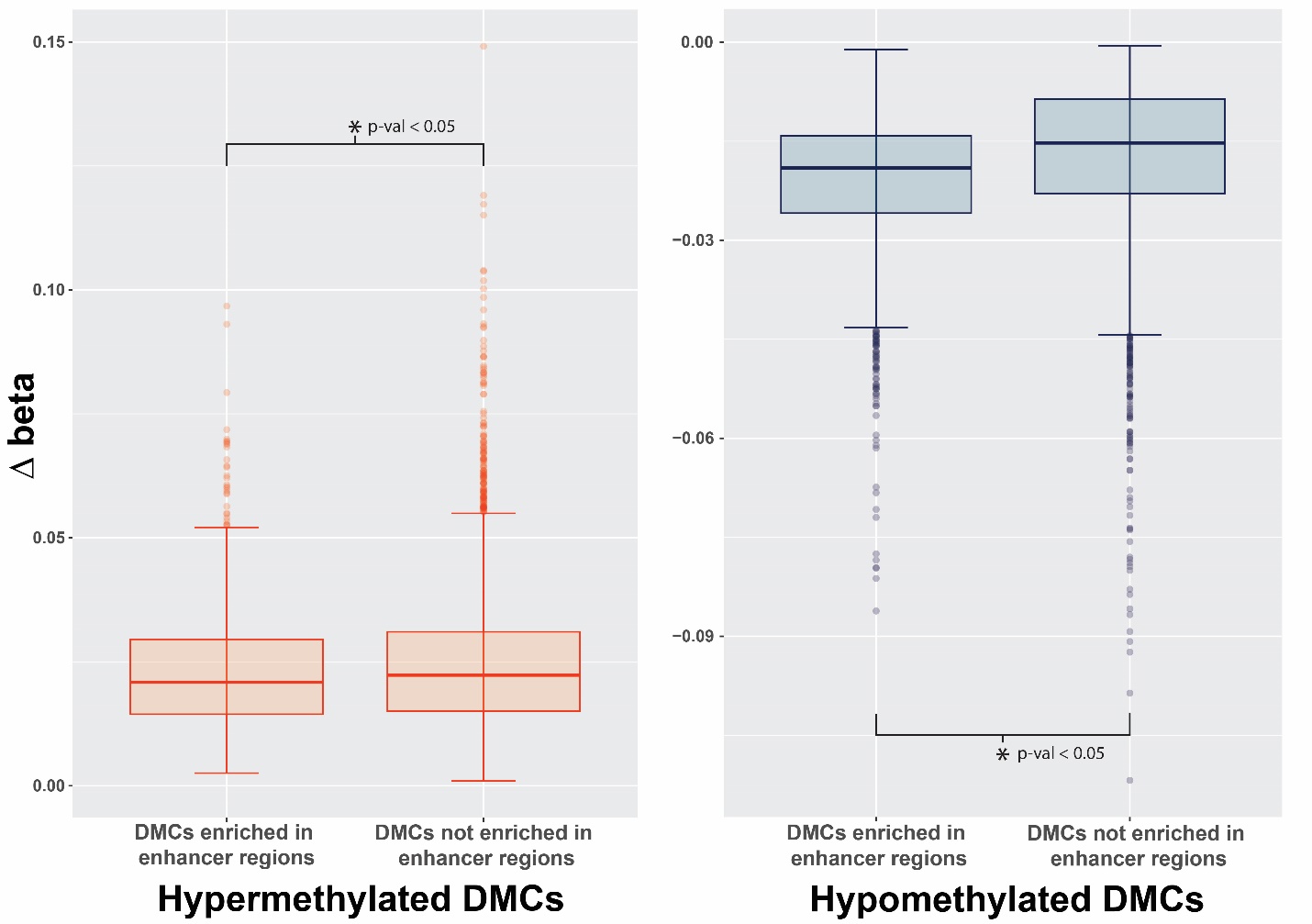

**Comparison of delta beta values at significant DMCs.** Analysis of the differences in beta values at DMCs from Sertoli, Granulosa, and iPSCs that were enriched at enhancer regions and associated with closer proximity to ERE elements or DMCs that were not enriched at enhancers that had a lower frequency of ERE elements in close proximity. Box plots display a high degree of similarity in the delta beta intensities measured for these specific DMCs in BPS-treated samples vs control samples. Interestingly, the differences between the distribution of beta values were sufficient to be significant based on the two-sample Kolmogorov-Smirnov test. These observed differences indicate that there is higher variability of the delta betas associated with hypomethylated changes occurring at DMCs associated with enhancers but not hypermethylation indicating a trend for a higher proportion of cells to have hypomethylated changes at these specific regions.
