## Supplementary file 8 for "Endocrine disruptor-induced epimutagenesis *in vitro*: Insight into molecular mechanisms"

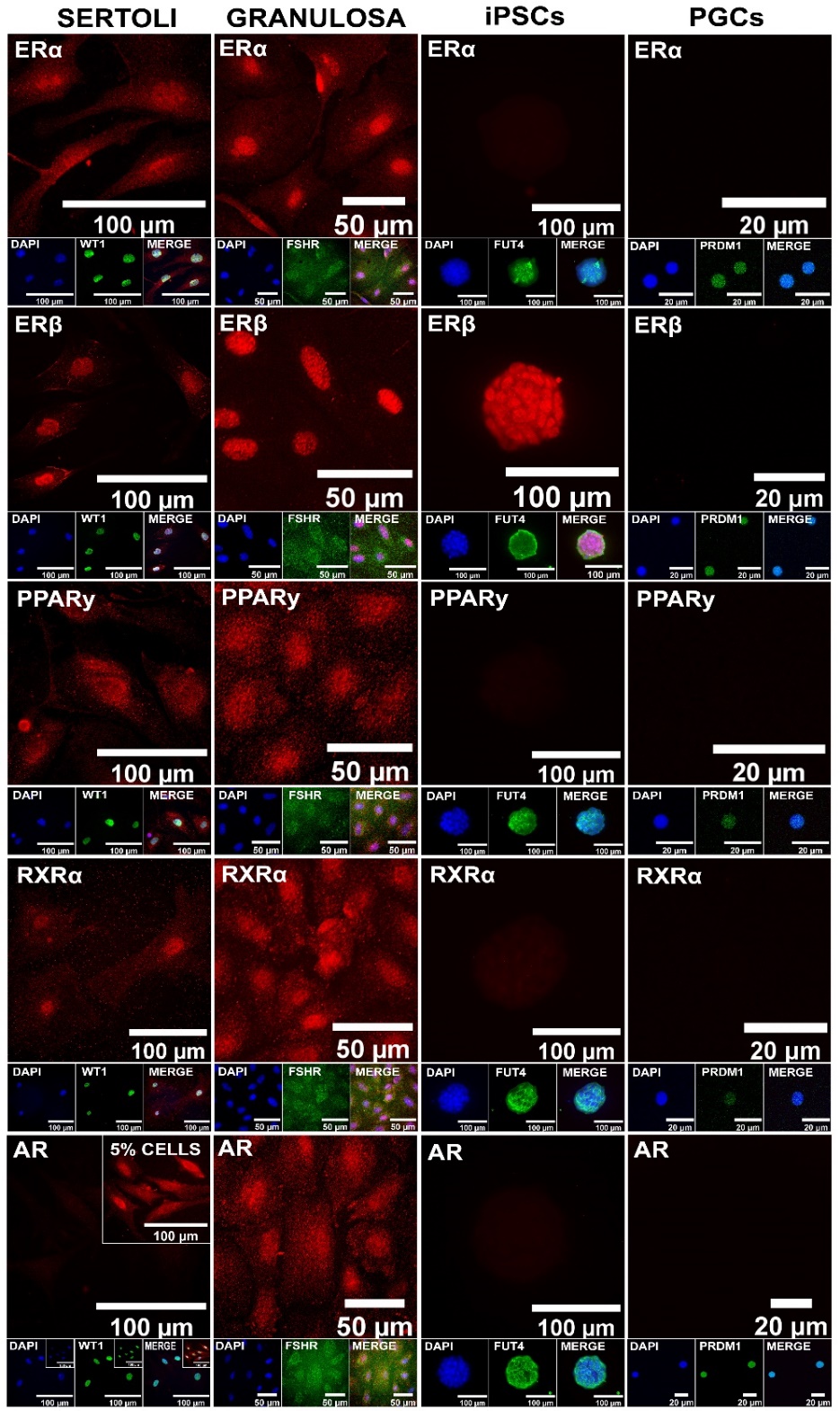

**Assessment of expression of additional endocrine receptors potentially involved in cell type-specific responses to BPS exposure.** Immunocytochemistry staining of expression of ERα, ERβ, PPARγ, RXRα, and AR is shown, along with staining for known cell type-specific markers. Somatic cell types express all receptors, pluripotent cells express ERβ but not ERα, PPARγ, RXRα or AR, and germ cells do not express any of the endocrine receptors.
