## Supplementary file 9 for "Endocrine disruptor-induced epimutagenesis *in vitro*: Insight into molecular mechanisms"

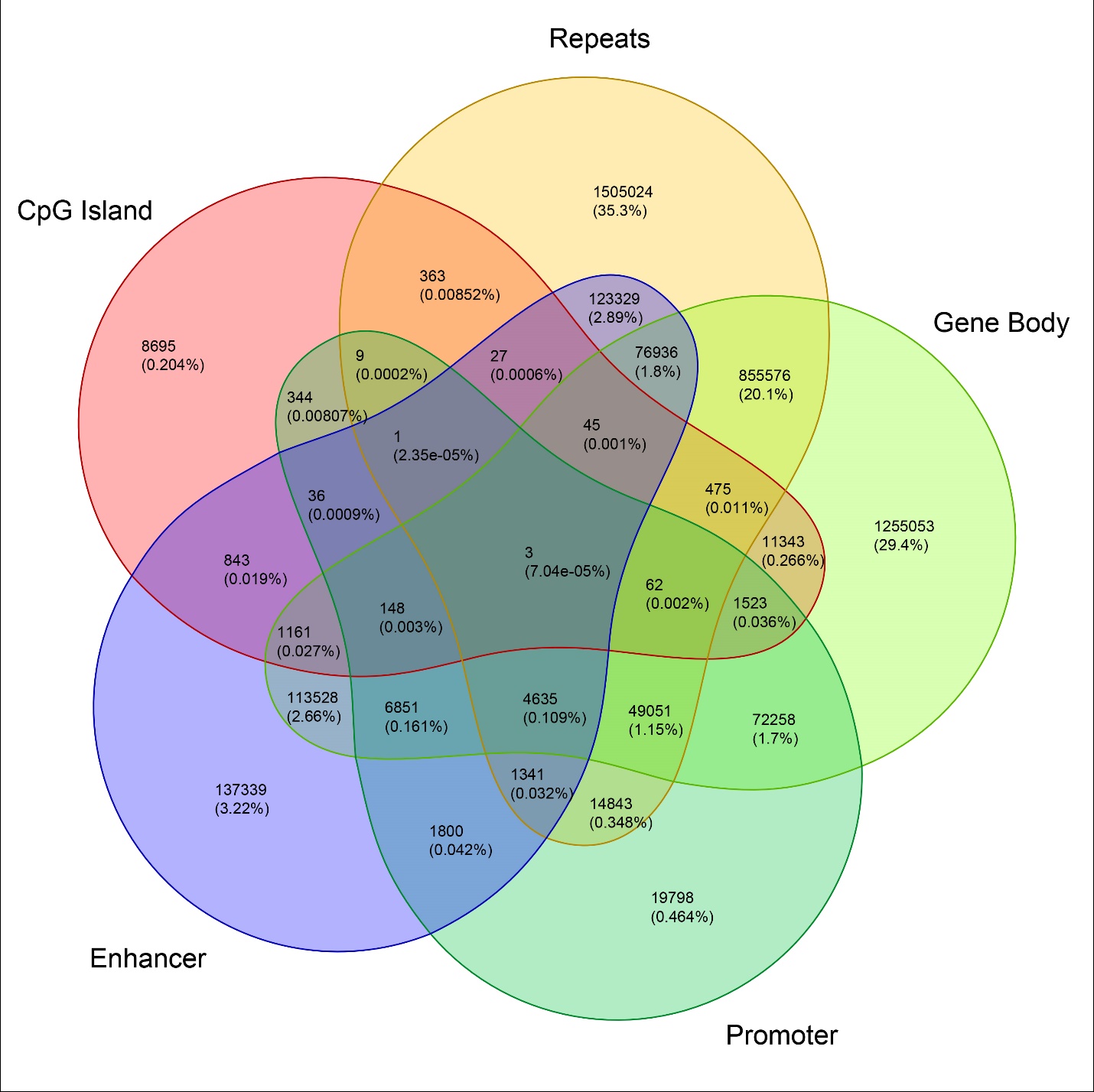

**Genome-wide annotation of ERE half-sites.** Venn diagram displaying the identification of ERE half-sites localized in known genic and intergenic regions.
