## Supplementary file 11 for "Endocrine disruptor-induced epimutagenesis *in vitro*: Insight into molecular mechanisms"

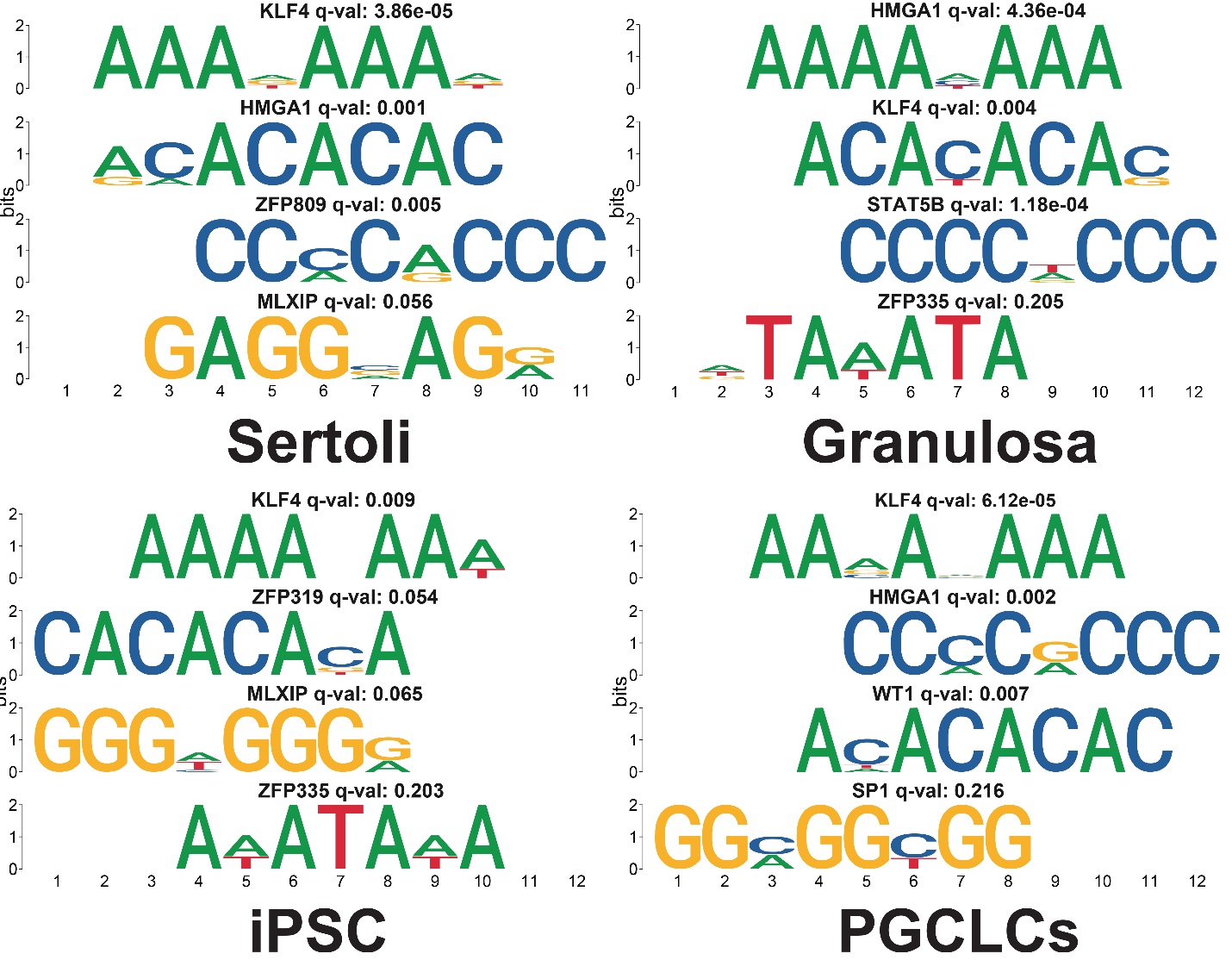

**Motifs near enriched DMCs.** Identification of the top 4 motif sequences (e-value < 0.05) within 500 bp of cell type-specific enriched DMCs that were either associated with enhancer regions in Sertoli, granulosa, and iPS cells or with transcription factor binding sites in PGCLCs. Each motif was compared with the JASPAR database for potential transcription factor binding capability associated with the motif. Transcription factors with potential binding capability are listed above each corresponding motif along with the adjusted p-value (q-value) of the association. Interestingly we see that the two most common motifs across all cell types was associated with either the chromatin remodeling transcription factor HMG1A or the pluripotency factor KLF4.
