## Supplementary file 12 for "Endocrine disruptor-induced epimutagenesis *in vitro*: Insight into molecular mechanisms"

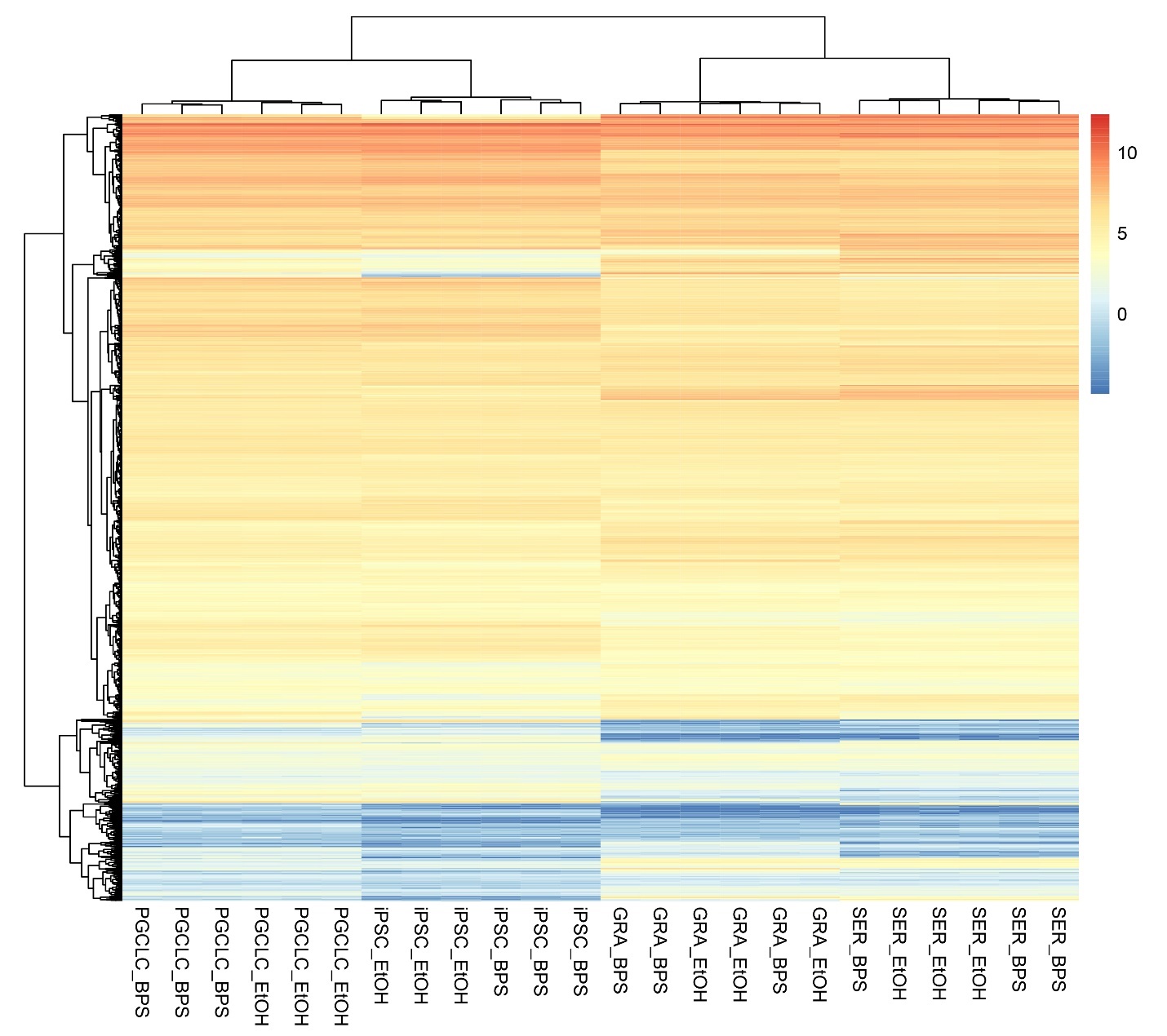

**Differential expression of potential endocrine-signaling independent DMCs.** Heatmap displaying the relative expression of genes with promoters enriched for apparent endocrine-signaling independent DMCs. The majority of these genes displayed a similar pattern of active expression in all cell types examined.
