## Supplementary file 14 for "Endocrine disruptor-induced epimutagenesis *in vitro*: Insight into molecular mechanisms"

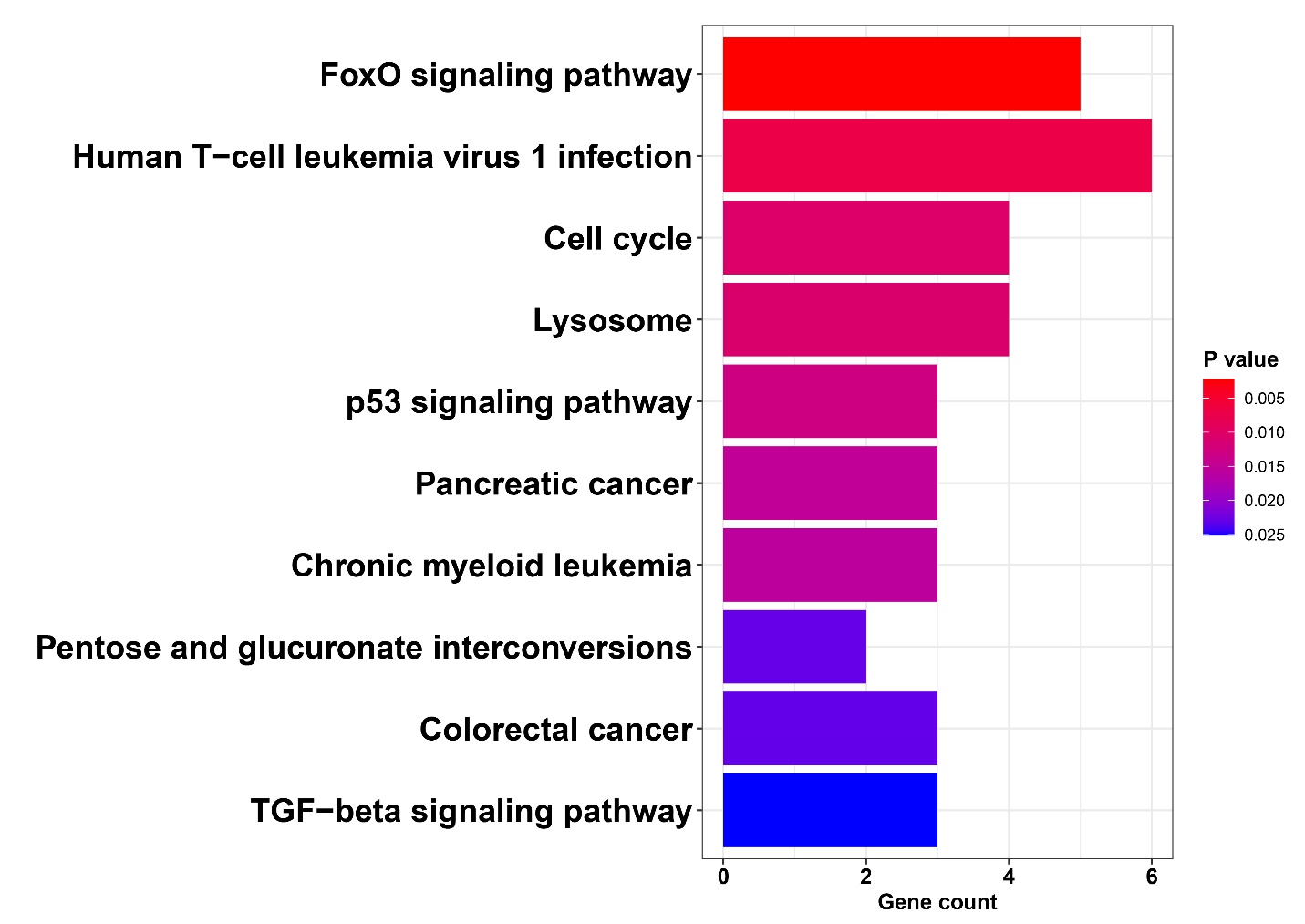

**KEGG pathway analysis of DEGs detected in both iPSCs exposed to BPS and PGCLCs derived from the exposed iPSCs.** Analysis of KEGG pathways associated with 138 BPS-induced DEGs that persisted during the transition in cell fate from BPS-exposed iPSCs to PGCLCs revealed genes primarily involved with cell cycle and apoptosis pathways.
