## Supplementary file 15 for "Endocrine disruptor-induced epimutagenesis *in vitro*: Insight into molecular mechanisms"

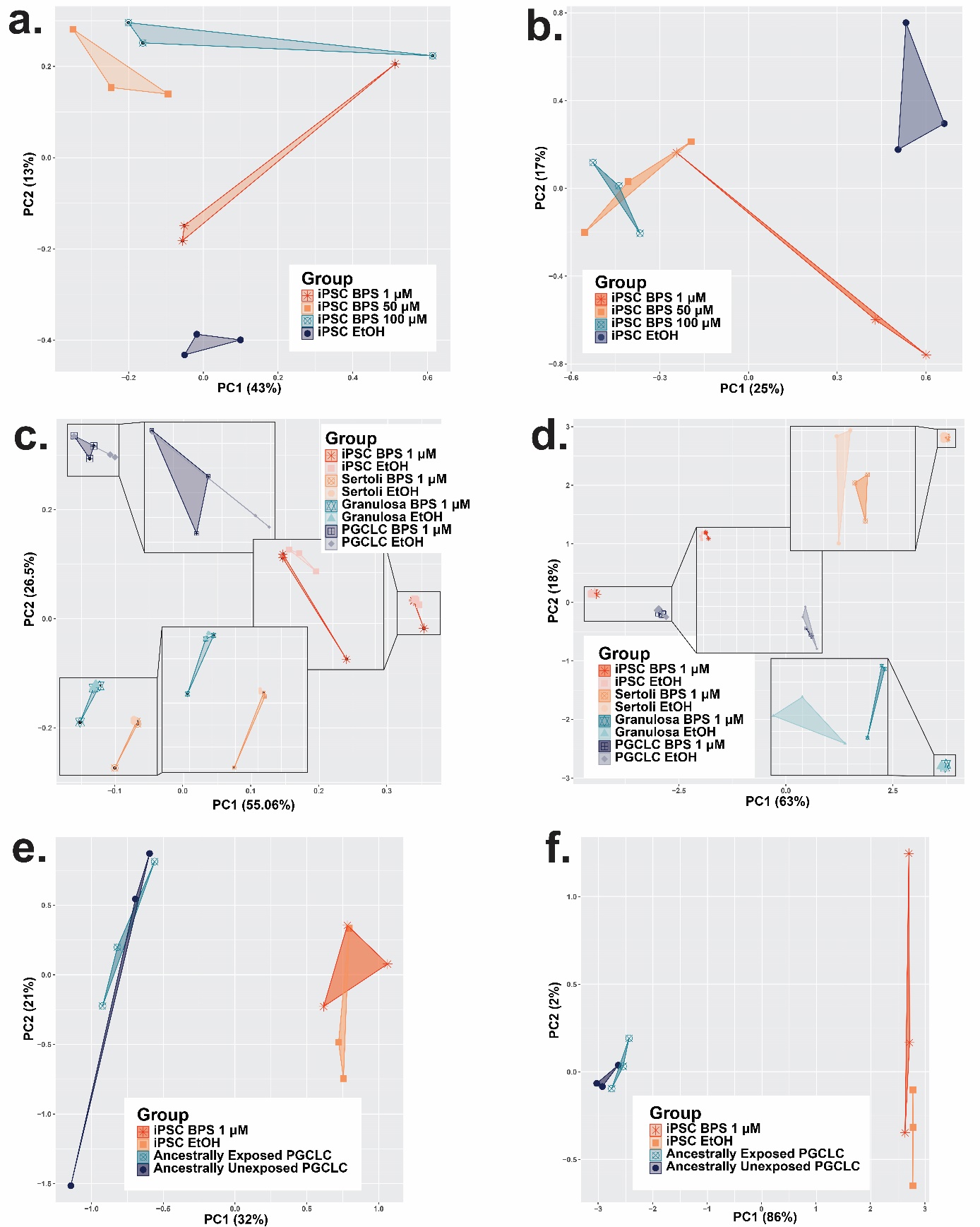
­

**Consistency among replicates and variation between experimental and control groups by assay.** Plots displaying principle component analyses of data from the **a-b)** BPS dose determination experiments showing dose-dependent susceptibility to BPS in iPSCs utilizing data from **a)** DNA methylation data from Infinium Beadchip array analysis and **b)** gene expression data from bulk RNA-seq analysis; **c-d)** cell type-specific susceptibility to BPS via changes in **c)** DNA methylation from DNA methylation Infinium Beadchip array data and **d)** gene expression from bulk RNA-seq data, and **e-f)** persistence of epimutations through transitions in cell states based on **e)** DNA methylation from EM-seq­ data and **f)** gene expression from bulk RNA-seq data. The dimensional reduction of the variation in plots **a-b)** demonstrates a partially additive dose-dependent relationship between the concentration of BPS added to the media and an increasing distinction between the control and treated samples, although this relationship plateaued at higher doses of BPS. In all other experiments, replicate samples clustered into regions based on distinct cell identity profiles with only minimal overlap between treatment and control samples. However, in plots e-f there is a lack of strong separation between treatment conditions in the second principle component. While the differences in all experiments were sufficient to produce DMCs/DMRs/DEGs this lack of separation displayed by the PCA could likely be increased by a larger sample size and indicates a limitation of only having triplicate replicates for this study.
