## Supplementary file 16 for "Endocrine disruptor-induced epimutagenesis *in vitro*: Insight into molecular mechanisms"

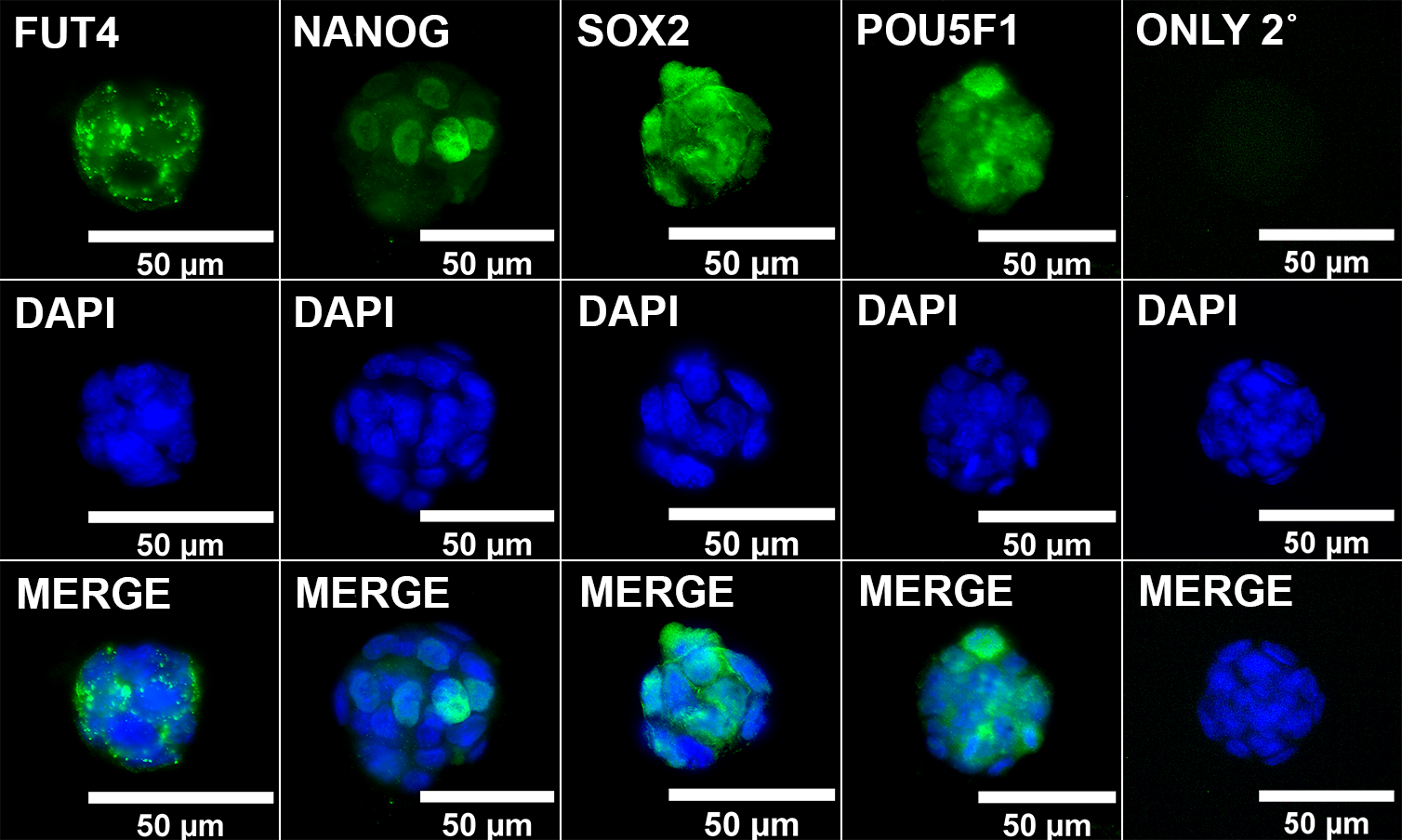

**ICC validation of MF5-9-1 iPSCs.** iPSCs from reprogrammed MEFs were validated for immunolabeling with known pluripotency markers along with negative control results when labeling was conducted with the secondary antibody only (ONLY 2˚).
