## Supplementary file 17 for "Endocrine disruptor-induced epimutagenesis *in vitro*: Insight into molecular mechanisms"

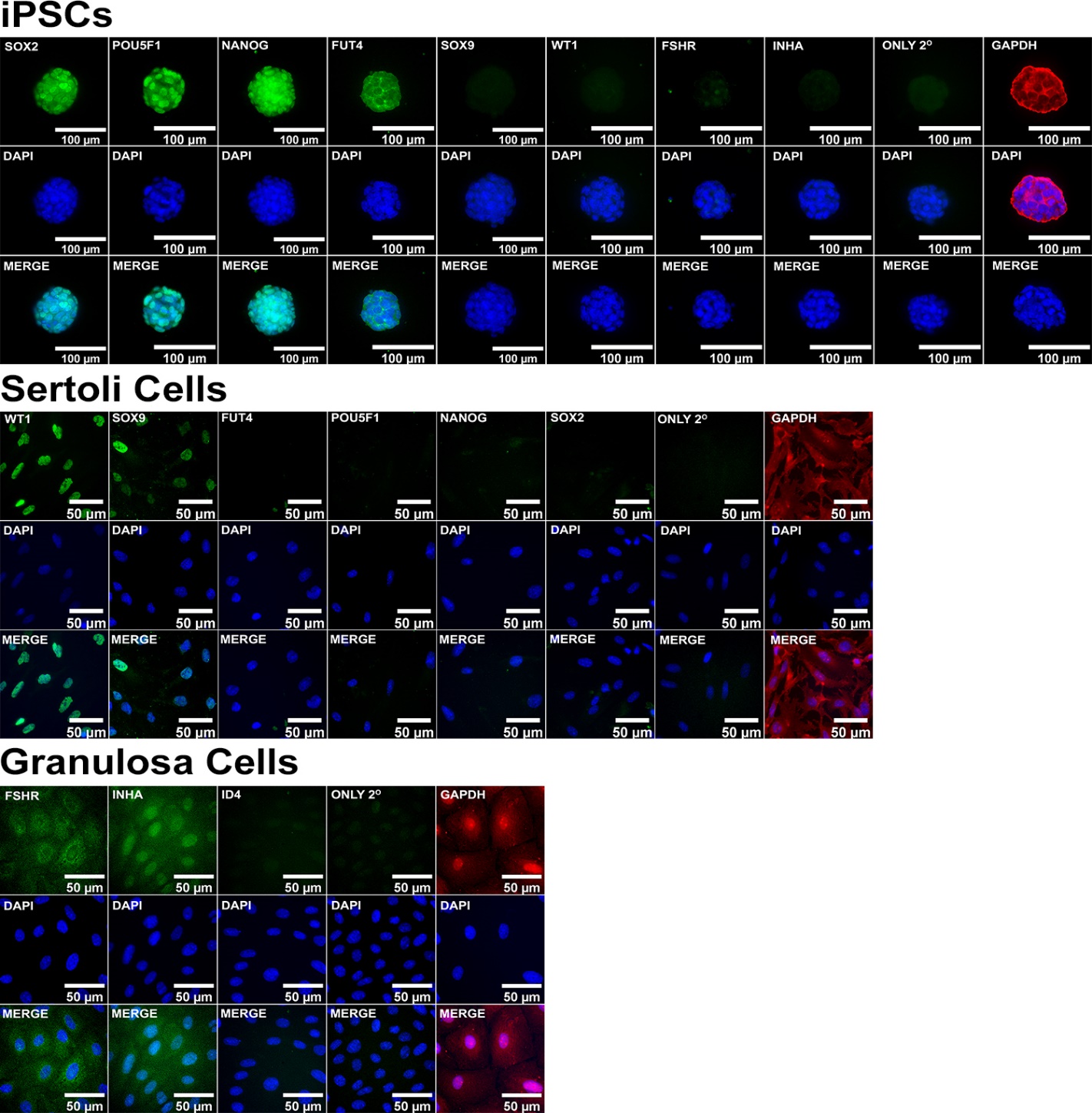

**ICC control staining of cell type-specific markers.** Validation of immunolabeling for known pluripotent and somatic cell type markers in iPSCs, Sertoli Cells, and Granulosa cells along with negative secondary antibody only (ONLY 2˚) and positive (GAPDH) controls.
