## Supplementary file 18 for "Endocrine disruptor-induced epimutagenesis *in vitro*: Insight into molecular mechanisms"

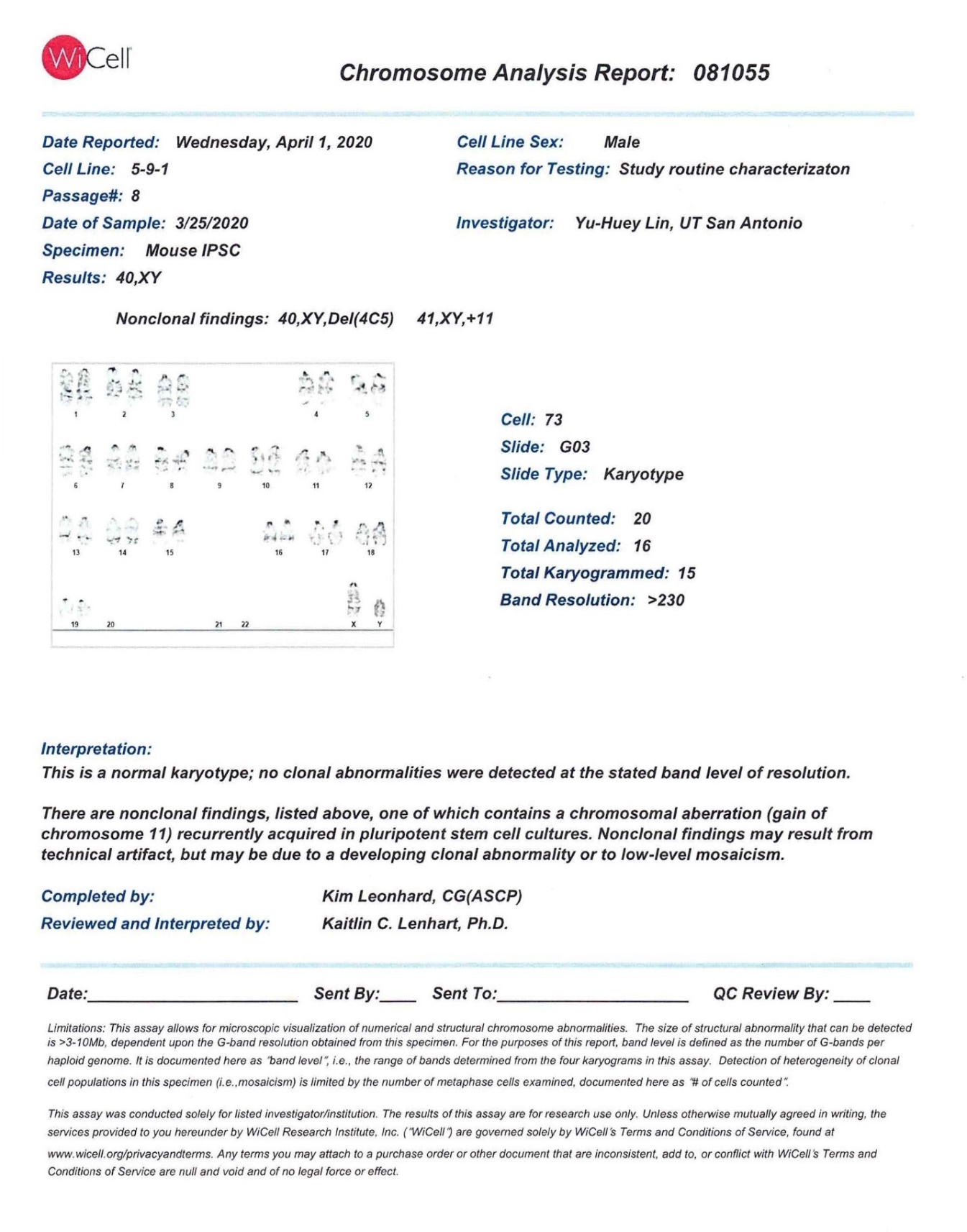

**Validation of normal karyotype analysis of MF5-9-1 iPSCs.** iPSCs from reprogrammed MEFs were validated for a normal karyotype prior to use in this project.
