## Supplementary file 19 for "Endocrine disruptor-induced epimutagenesis *in vitro*: Insight into molecular mechanisms"

**iPSC, EpiLC, and PGCLC Culture Media Components**

Stock solutions to prepare before making media.

1. Insulin stock sol. (25 mg/mL), (Sigma I-1882):

100 mg in 4 mL (0.01 M HCl), overnight at 4C

aliquot 0.5 mL (total 8 tubes) 🡪 -20^o^C

2. Apotransferrin stock sol. (100 mg/mL), (Sigma T-1147):

500 mg in 5 mL (water), overnight at 4 C

aliquot 0.5 mL (10 tubes) 🡪 -20^o^C

3. Progesterone stock sol. (0.6 mg/mL) (Sigma P8783):

10 mg in 1 mL ethanol (10 mg/mL)

🡪 60 μL of stock (10 mg/mL) + 940 μL ethanol 🡪 filter through a 0.22 μm filter

aliquot 100 μL (10 tubes) 🡪 -20^o^C

4. Putrescine stock sol. (160 mg/mL) (Sigma P5780):

320 mg in 2 mL water 🡪 0.22 filter

aliquot 60 μL (~30 tubes) 🡪 -20^o^C

5. Sodium selenite stock sol. (3mM) (Sigma S5261, 10G, MW:172.49)

300 mM: 51.88 mg in 1 mL water 🡪 0.22 filter

10 μL (300 mM stock) + 990 μL water = 3 mM

aliquot 50 μL (~20 tubes) 🡪 -20^o^C

6. Activin A stock sol. (50 μg/mL) (120-14, total 10 μg/vial, need more)

20 μg (2 vials) + 400 μL water 🡪 5 μL aliquot (~40 tubes) 🡪 -20^o^C

7. bFGF stock sol. (10 μg/mL) (13256-029, 10 μg/vial)

10 μg (total) in 1 mL (10 mM Tris, pH 7.6 with 0.1 % BSA)

🡪 aliquot 20 μL 🡪 -20^o^C

8. Recombinant human BMP4 stock sol. (50 ug/mL) (314-BP-010)

60 ug (total) + 1200 μL (4 mM HCl with 0.1% BSA)

🡪 aliquot 105 μL (12 tubes total) 🡪 -80^o^C

9. Stem Cell Factor (SCF) stock sol. (50 ug/μL) (455-MC-010)

10 ug (total) + 200 μL (PBS with 0.1% BSA)

🡪 aliquot 21 μL (10 tubes total) 🡪 -80^o^C

10. Recombinant mouse EGF stock sol. (500 ug/mL) (2028-EG-200)

200 ug (total) + 400 μL (PBS with 0.1% BSA)

🡪 aliquot 10 μL (60 tubes) 🡪 -80^o^C

**iPSC media**

**N2B27 media for culturing iPSCs in feeder-free conditions**

**N2 media components (5 mL)**

| Name | Volume | Catalog  Number | Storage Temp |
| --- | --- | --- | --- |
| DMEM/F12 | 3.6 mL | 21041-025 | (4^o^C) |
| Insulin stock sol. | 0.5 mL | I-1882 | (-20^o^C) |
| Apotransferrin stock sol. | 0.5 mL | T-1147 | (-20^o^C) |
| 7.5% BSA sol. | 330 μL | 15260-037 | (4^o^C) |
| Progesterone stock sol. | 16.5 μL | P8783 | (-20^o^C) |
| Putrescine stock sol. | 50 μL | P5780 | (-20^o^C) |
| Sodium selenite stock sol. | 5 μL | S5261 | (-20^o^C) |

**N2B27 media components (1 L)**

| Name | Volume | Catalog  Number | Storage Temp |
| --- | --- | --- | --- |
| N2 | 5 mL | Table 1 | (RT) |
| DMEM/F12 | 495 mL | 21041-025 | (4^o^C) |
| Neurobasal | 480 mL | 12348-017 | (4^o^C) |
| B27 (50X) | 10 mL | 12587-010 | (-20^o^C) |
| Pen/Strep | 5 mL | 15070063 | (-20^o^C) |
| L-glutamine stock sol. | 5 mL | 35050061/Glutamax | (4^o^C) |
| β-ME stock sol., 50 mM | 1.8 mL | 21985-023 | (4^o^C) |

Make a 40 mL aliquot of N2B27 and store at -80^o^C.

**N2B27 2i-LIF media components (40 mL)**

| Name | Volume | Catalog  Number | Storage Temp |
| --- | --- | --- | --- |
| N2B27 | 40 mL | Table 2 | (-80^o^C) |
| CHIR99021 | 12 μL | 1677-5 | (-80^o^C) |
| PD0325901 | 3.2 μL | 04-0006-02 | (-80^o^C) |
| ESGRO | 4 μL | ESG1107 | (4^o^C) |

Store at 4^o^C for up to 2 weeks.

**EpiLC differentiation**

Plate coating solution.

1. Human plasma fibronectin (1 mg/mL, store at 4C) (1 mg/mL, FC010, 1 mg total = 1 mL),

Working solution: For a 12-well plate: dilute 10 μL fibronectin with 600 μL PBS to obtain 16.6 μL/mL fibronectin solution. Pipet the solution into one well of a 12-well plate, incubate for at least 1hr at 37^o^C and then aspirate the fibronectin solution just before plating iPSCs.

**N2 EpiLC differentiation (~5 mL)**

| Name | Volume | Catalog  Number | Storage Temp |
| --- | --- | --- | --- |
| N2B27 | 5 mL | Table 2 | (-80^o^C) |
| Activin A stock sol. | 2 μL | 120-14 | (-20^o^C) |
| bFGF stock sol. | 6 μL | 13256-029 | (-20^o^C) |
| KSR | 50 μL | 10828-028 | (-20^o^C) |

**PGCLC differentiation**

**GK15 (10 mL)**

| Name | Volume | Catalog  Number | Storage Temp |
| --- | --- | --- | --- |
| GMEM | 8.082 mL | 11710-035 | (4^o^C) |
| KSR | 1.5 mL | 10828-028 | (-20^o^C) |
| Non-EAA | 100 μL | 11140-050 | (4^o^C) |
| Sodium pyruvate | 100 μL | 11360070 | (4^o^C) |
| Pen/Strep | 100 μL | 15070 | (4^o^C) |
| L-glutamine | 100 μL | 35050061/Glutamax | (4^o^C) |
| β-ME, 50 mM | 18 μL | 21985-023 | (4^o^C) |

**PGCLC differentiation media (10 mL)**

| Name | Volume | Catalog  Number | Storage Temp |
| --- | --- | --- | --- |
| GK15 | 9.878 mL | Table 5 | (4^o^C) |
| BMP4 | 100 μL | 314-BP-010 | (-80^o^C) |
| SCF | 20 μL | 455-MC-010 | (-80^o^C) |
| LIF | 1 μL | ESG1107 | (-80^o^C) |
| EGF | 1 μL | 2028-EG-200 | (-80^o^C) |

**TrypLE wash media (75 mL)**

| Name | Volume | Catalog  Number | Storage Temp |
| --- | --- | --- | --- |
| 7.5% BSA sol. | 1 mL | 15260-037 | (4^o^C) |
| DMEM/F12 | 74 mL | 21041-025 | (4^o^C) |
