## Supplementary file 20 for "Endocrine disruptor-induced epimutagenesis *in vitro*: Insight into molecular mechanisms"

**Preparation of primary cultures of Sertoli cells from mice**

**Materials:**

500 mL DPBS	(Thermo Fisher: Cat. No. 14040133)

DNaseI. 6.64 mg/mL in DPBS	(Sigma: Cat. No. DN-25)

Trypsin 2.5% (10X Gibco)	(Gibco-BRL: Cat. No. 15090-046)

Soybean trypsin inhibitor, 3.4 mg/mL in DPBS	(Gibco-BRL: Cat. No. 17075-011)

Collagenase type IV, 0.70 mg/mL	(Worthinton: Cat. No. LS004188)

Sertoli cell media containing retinoic acid	(ScienCell^TM^: Cat. No. 4521)

**Equipment:**

Pyrex Trypsinizing Flask 300 mL Standard Flask (Fisher Cat. No. 09-552-45)

**Procedure:** (maintain sterility throughout procedure)

A. **Preparation of testes tissue**
1. Sacrifice mice by CO_2_.
2. Wipe down abdomen with 70% EtOH.
3. Make abdominal incision.
4. Remove the testes and immerse in DPBS in a 50 mL conical tube.
5. Remove tubule bundle from tunica.
a. Place testes on parafilm, grasp at one end, and make a small incision in the tunica at the opposite end.
b. Push the tubules out of tunica by using the side of closed scissors.
c. Weigh flask with fresh DPBS (this is the DPBS volume from the first step below so you must estimate for now (W_1_): g
d. Place tubules in DPBS and weigh (W_2_): g
e. Calculate testes weight (W_T_) = (W_2_ - W_1_): g.
B. **Preparation of Sertoli cells**
1. Prepare Erlenmeyer flask by adding:
a. Trypsin, 10x (2.5 x W_T_) =	mL
b. DPBS (22.5 x W_T_) =	mL
c. DNaseI (100 x W_T_)= μL (Start with 100 μL of DNAse1 and then added 1 mL at the 20 min mark and another 1 mL at the 25 min mark.)
   DNaseI is very critical, always favor more rather than less and be certain about the concentration when preparing the stock. This along with the flask could be why you are having clumping problems.

Before adding tissue place in a clean petri dish and using a clean razor blade chop downward across the tissue and then rotate the plate 90^o^ and chop across the tissue again until the tissue loses its grape structure as a pre-processing step to help break up the tubules so they digest easily.

Add tissue to the flask. Cover flask with foil, incubate at 37^o^C for 25-30 min, with occasional swirling. During the last ten minutes, tissue should come apart rapidly. If after 20 min strings seen while swirling (this is very empirical when properly digesting the tissue will break up into pieces approx 2-3 mm in length and be independent of each other not clumped together), add DNaseI at (50 x W_T_	μL), after an additional 5 min add DNaseI at (25 x W_T_	μL).

2. When tubules are separated, move flask to sterile hood, let tubules settle for 5 mins. Then remove and discard overlying liquid. The solution will be very sticky and the tissue will want to come out with the supernatant. Don’t lose too many cells here, if you can not get all the supernatant off just go ahead and add the trypsin inhibitor, but use another 0.5 mL on top of what you have figured to compensate for the increased volume.
1. Add trypsin inhibitor (2.5 x W_T_)=	mL. Swirl and incubate for 1 min.
2. Add (25 x W_T_) = mL fresh DPBS, swirl, allow to settle. (Wait to add this if you did not get much of the supernatant off before, too much volume will increase the buoyancy and not let the cells settle out.
3. Remove supernatant, add (20 x W_T_)=	mL DPBS, swirl,

allow to settle.

4. Remove supernatant, add

a. (7.5 x W_T_)=	mL DPBS

b. Collagenase (2.5 x W_T_)=	mL

c. DNaseI (25 x W_T_)=	μL (Add in 1mL here of DNaseI instead if below 1 mL)

5. Incubate in water bath at 37^o^C for 10 min. After 5 min, swirl continuously. (During the last 3 min aliquots are checked by microscope to monitor the release of peritubular cells. A drop of suspension is placed on a glass slide and the tissue is viewed under a coverslip at 10x mag. The coverslip is moved slightly and as the tissue moves peritubular cells can be seen to peel off. The tissue is very fragile at this stage and must not be allowed to over digest. Digestion is complete when the surface of large fragments no longer appears smooth.
6. Transfer tubules to 50 mL conical tube, wash the flask with DPBS, and centrifuge at 200xg for 2 min.
7. Remove supernatant, add 30-35 mL DPBS, and centrifuge again.
8. Repeat step 9.
9. Remove supernatant and resuspend and plate cells in Sertoli cell media containing retinoic acid from ScienCell^TM^.
10. Cells are incubated at 37^o^C in 5% CO_2_, in a humidified chamber. Medium is changed every 48 hours after 24-36 hours shock the cells with 10 mM Tris-HCl, pH 7.4 for 2 min. This will remove most of the contaminating germ cells. Make this media by dissolving 12.1g of Trisbase in 80 mL of ddH_2_O and adjust the pH to 7.4 with HCl. This will make a 1M stock. To make a 10 mM solution, add 1mL of this 1M stock to 99 mL of ddH_2_O and filter sterilize.
