## Supplementary file 21 for "Endocrine disruptor-induced epimutagenesis *in vitro*: Insight into molecular mechanisms"

**Preparation of primary cultures of granulosa cells from mice protocol**

**Materials:**

70% EtOH	(Fisher: Cat. No. BP8203)

500 mL DPBS	(Thermo Fisher: Cat. No. 14040133)

BSA	(Sigma: Cat. No. A9085)

Collagenase type IV	(Worthinton: Cat. No. LS004188)

Glass Pipettes	(Fisher: Cat. No. 13-678-20A)

DMEM/F12	(Gibco Cat. No. 21041-025)

FBS	(Gibco Cat. No. 10082-147)

Penicillin/Streptomycin 100X	(Gibco Cat. No. 15140-122)

0.25% Trypsin-EDTA	(Gibco Cat. No. 25200-072)

**Procedure:**

A. **Preparation of media**
1. Prepare 50 mL of DPBS with 0.5% BSA by dissolving 0.25 g of lyophilized BSA powder into 50 ml of DPBS medium.
2. Collagenase type IV solution (540 U/pair of ovaries) in 3 mL of DPBS.
3. Prepare a glass pipette with a pointed tip that you can use to transfer the follicles. Set up an alcohol flame. Holding a glass pipette from both ends place the thicker section of glass just before the pipette tapers off over the hottest part of the flame. Rotate the tubing to make sure it is evenly heated. Continue to heat the glass until it starts to melt and becomes flexible. Then, with a quick motion, pull the glass so it forms a thin tube section. This will take some trial-and-error force that will give you the size that works best. Etch the glass and tap near the etched section to make a clean break in the tube. Finally, sterilize the tube and remove jagged edges by quickly running the completed tube through the flame. Attach the glass pipette to some form of controlled suctioning device (i.e. tubing connected to small syringe)
4. Prepare 50 mL of DMEM/F12 with 15% FBS by adding 7.5 mL of FBS and 0.25 mL of 100X Penicillin/Streptomycin stock solution to 42.25 mL of DMEM/F12 medium.
B. **Isolation of ovarian follicles from mice**
1. Sacrifice mice by CO_2_.
2. Wipe down abdomen with 70% EtOH.
3. Make abdominal incision.
4. Remove the ovaries and immerse in DPBS in petri dish.
5. Decap the ovaries by resecting away fat and connective membranes that surrounds and protect the ovary.
6. Transfer the decapped ovaries to a clean petri dish with DPBS and no BSA. This will help to clean off any contaminating hairs, or fat and blood droplets released while decapping the ovaries.
7. Section the ovaries into fourths by cutting them with the needle and forceps. This will help the collagenase digest the tissue and release follicles in the next step.
8. After the ovaries have been divided into fragments add them to the activated collagenase digest solution and let the ovaries digest for 25 mins under constant agitation.
9. At the end of 25 mins add in DPBS containing 0.5% BSA to inactivate the collagenase then centrifuge the tube at 60xg for 5 mins to pellet the follicles^214^.
10. After the tissue digest has finished spinning remove the supernatant and resuspend in 7 mL of DPBS with 0.5% BSA and transfer to a new petri dish to individually pick out follicles.
B. **Preparation of granulosa cells**
1. Take the petri dish containing resuspended ovarian tissue and place it under stereomicroscope.
2. Move the preantral follicles over to the clean droplet of DPBS with 0.5% BSA. It’s best to move only a few follicles over at a time as pulling up many follicles can cause them to sediment and stick to the inside of the glass pipette if they are pulled up too far inside.
3. Once you have moved over enough follicles, set up a 1.5 mL microcentrifuge tube containing 1 mL of 0.25% activated trypsin.
4. Gently swirl the dish containing the preantral follicles to get them to clump together in the center and collect them with a 1000 μL micropipette before transferring them to the droplet containing the trypsin EDTA.
5. Let the follicles digest for 5 minutes. Pipette the digesting cells to help break up the follicles during the incubation period.
6. Spin tube at 1000xg for 5 mins to pellet the granulosa and oocyte cells.
7. Remove the supernatant and resuspend pellet in DMEM/F12 supplemented with 15% FBS and plate cells.
