## Supplementary file 22 for "Endocrine disruptor-induced epimutagenesis *in vitro*: Insight into molecular mechanisms"

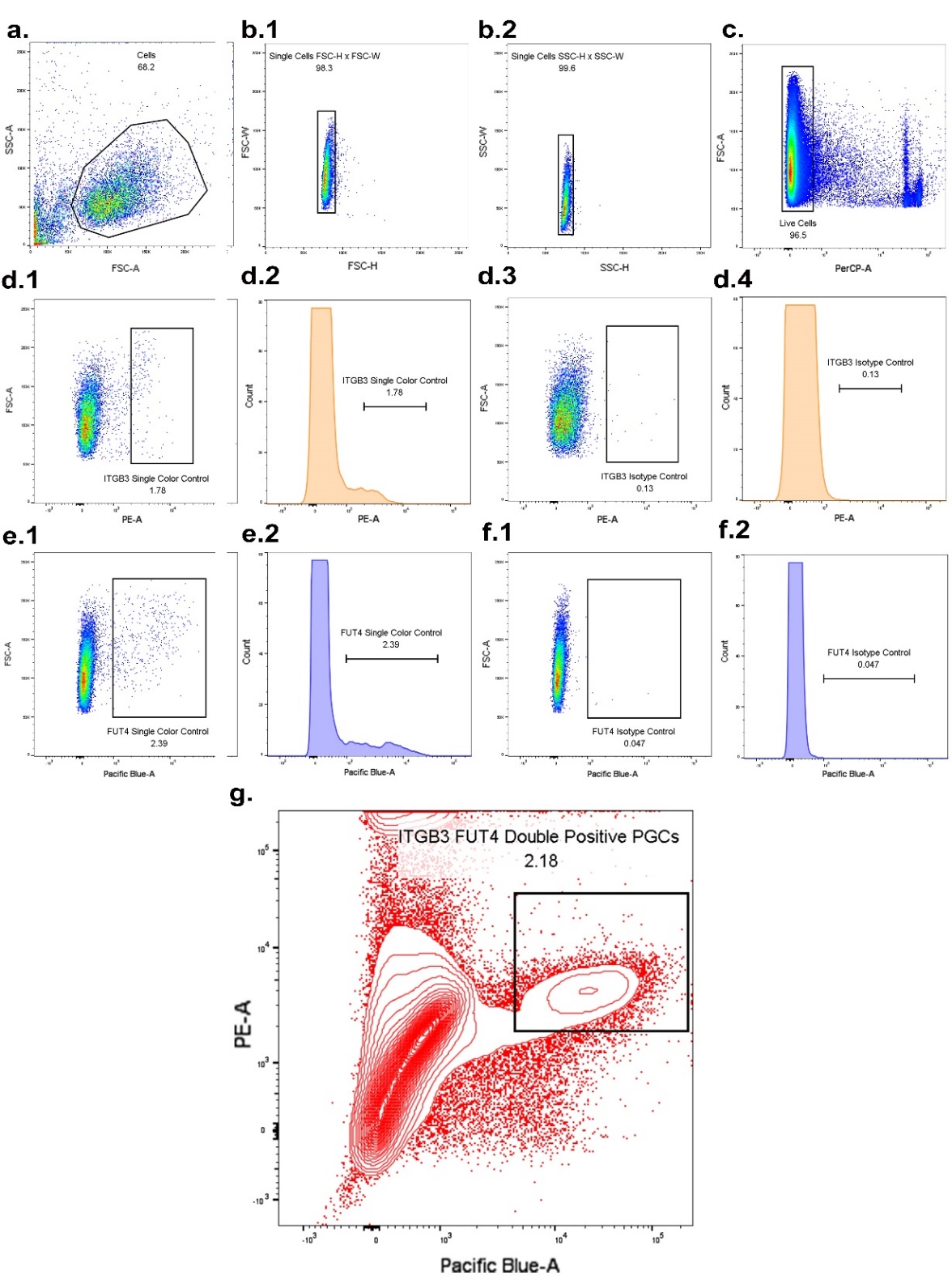

**FACS sorting for ITGB3/FUT4 enriched primordial germ-cell like cells. a)** Gating for cells. **b.1-2)** Gating for singlet cells. **c)** Gating for live cells. **d.1-2)** Single color control ITGB3 positive cells. **d.3-4)** IgG isotype control. **e.1-2)** Single color control FUT4 positive cells. **f.3-4)** IgG isotype control. **g)** Sorting for PGCLC-enriched ITGB3/FUT4 double positive population (2.18% of total cells).
