## Supplementary file 23 for "Endocrine disruptor-induced epimutagenesis *in vitro*: Insight into molecular mechanisms"

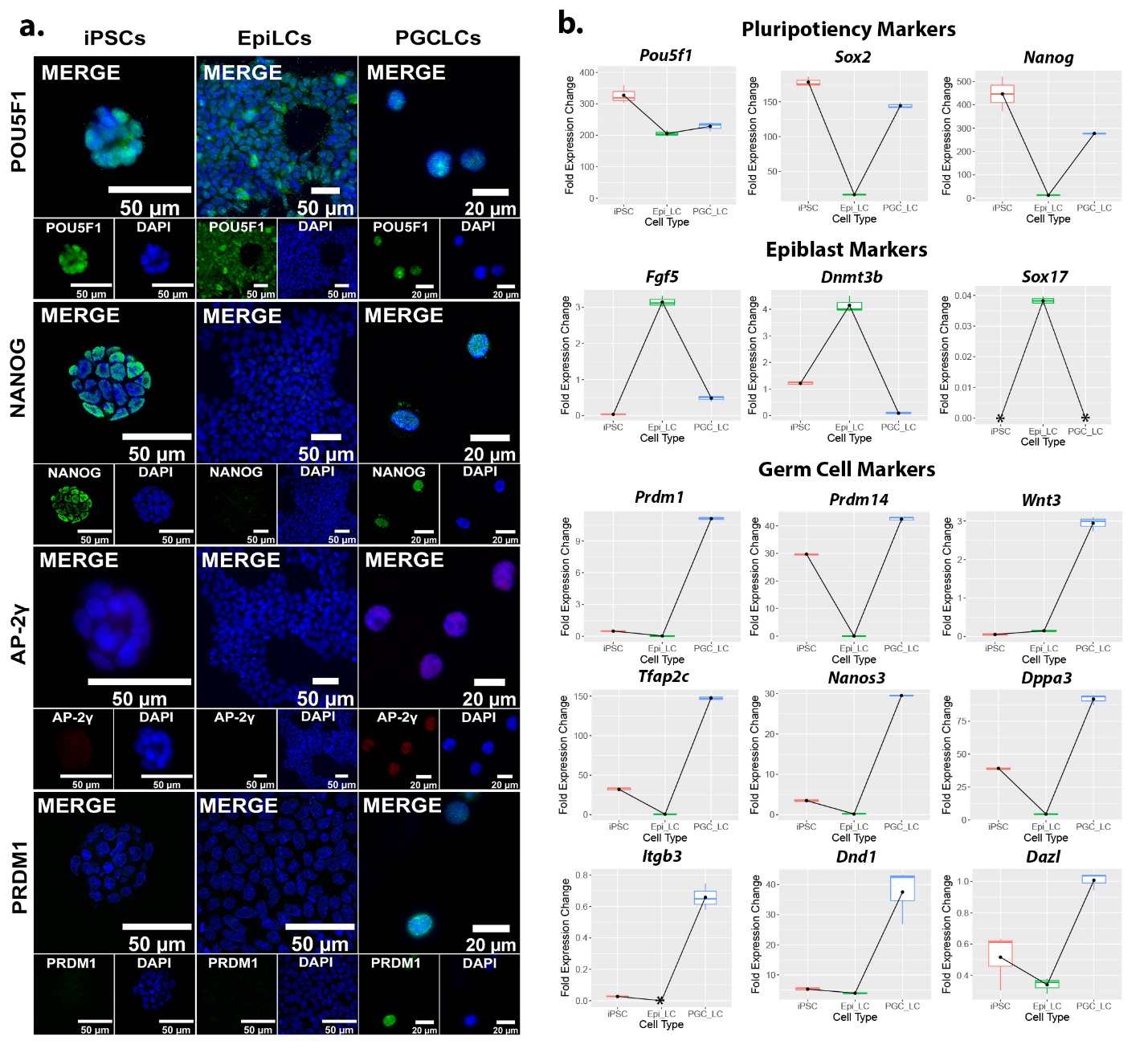

**Relative expression of markers for PGCLC induction from iPSCs. a)** ICC of pluripotency and germ cell marker expression throughout the transition from iPSCs to PGCLCs. **b)** qRT-PCR of pluripotency, epiblast, & germ cell markers indicating gene expression profiles during induction of PGCLCs from iPSCs. Each gene fold expression is relative to the housekeeping gene *Gusb* using the ∆Cq method. The symbol * indicates that the expression of transcripts in the sample was either non-existent or so low as to be undetectable by qRT-PCR.
