## Supplementary file 24 for "Endocrine disruptor-induced epimutagenesis *in vitro*: Insight into molecular mechanisms"

**qRT-PCR primers**

| Gene | Forward | Reverse | Efficiency |
| --- | --- | --- | --- |
| *Pou5f1* | TTCAGCCAGACCACCATCTGT | GGTTCTCATTGTTGTCGGCTTC | 106.02% |
| *Nanog* | GTGCTTGCTTGTCCTTGGAG | AGCTGGCATCGGTTCATCAT | 103.86% |
| *Tfap2c* | AGGCATCTCATTTCTGCGGG | CACGGACAGGCTTAGAGGTC | 101.37% |
| *Prdm1* | GAAAAATGGGAGCCCCGACA | AAGACGGAAGGGGACTGTGA | 92.60% |
| *Prdm14* | ACAGCCAAGCAATTTGCACTAC | TTACCTGGCATTTTCATTGCTC | 104.57% |
| *Sox2* | TTGGGAGGGGTGCAAAAAGA | TTCTAGTCGGCATCACGGTT | 98.20% |
| *Sox17* | CAAAGCGAAGATTGCAGGGT | TTTCCAGTCCCTGGTCAGTC | 107.78% |
| *Fgf5* | TACGTGGCCCTGAACAAGAG | TGGAACAGTGACGGTGAAGG | 108.72% |
| *Dnmt3b* | GCTTTGCTTCACTGGGTCTC | AACTGATGGGGTACTGACGC | 98.12% |
| *Wnt3* | CCTGGACCACATGCACCTAAA | CGGAGGCACTGTCGTACTTG | 95.11% |
| *Nanos3* | CACTACGGCCTAGGAGCTTGG | TGATCGCTGACAAGACTGTGG | 90.56% |
| *Dppa3* | TGAACGGGACAGTGAGCCA | AGCCAGGGCAGCGTACAAT | 99.12% |
| *Itgb3* | ACGGGACTTTTGAGTGTGGG | GCTGAGAGGGTCGGTAATCC | 100.54% |
| *Dnd1* | AATGGGTTAAGCAGAGCCTGT | AGGGCAAGGTTCCTCACAAC | 96.18% |
| *Dazl* | GCAGAGAACCTTTGTACCCAC | TAGCAAGGTGCTGTTCCAGTG | 102.75% |
| *Gusb* | GCAGCCCTTCGGGACTTTAT | TCCTCAACACCACTCTCATGTC | 97.17% |
