## Supplementary file 25 for "Endocrine disruptor-induced epimutagenesis *in vitro*: Insight into molecular mechanisms"

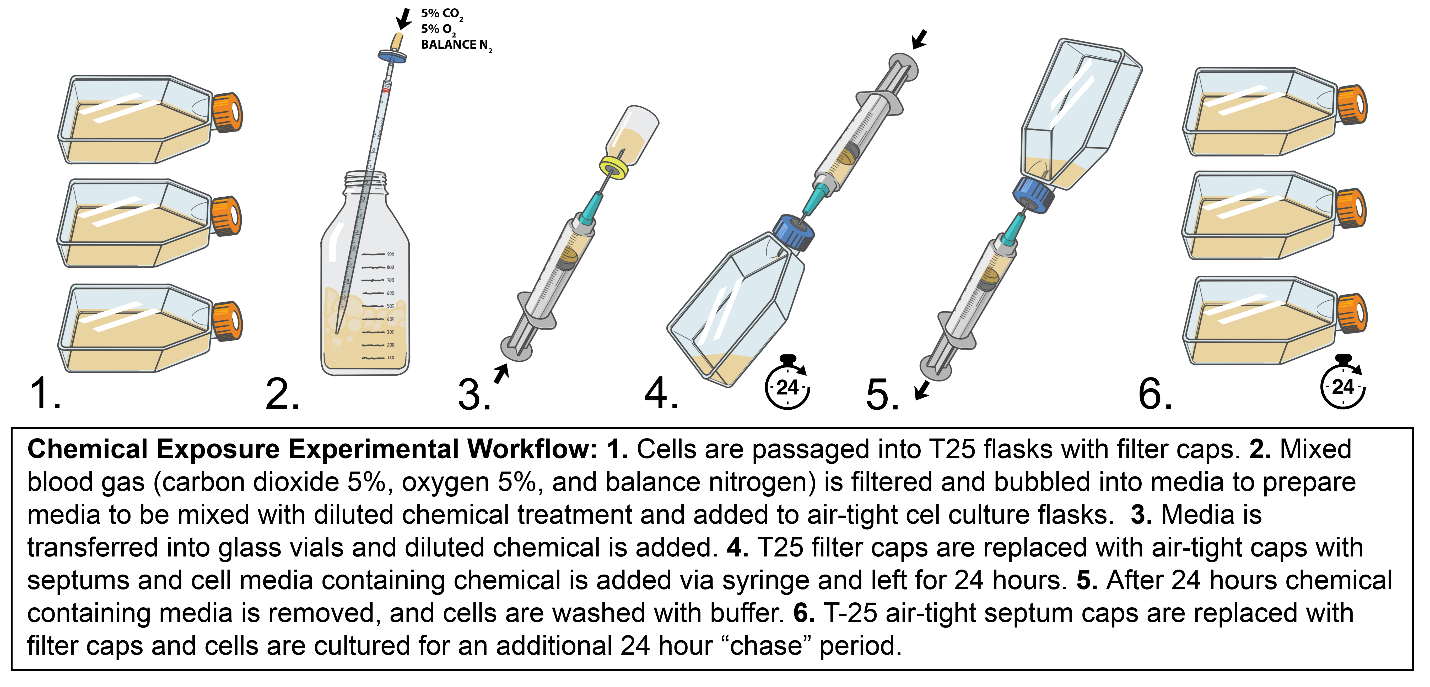

**Chemical exposure experimental workflow. 1)** Cells are passaged into T-25 flasks with filter caps. **2)** Mixed blood gas (carbon dioxide 5%, oxygen 5%, and balance nitrogen) is filtered and bubbled into media to prepare media to be mixed with diluted chemical treatment and added to air-tight cell culture flasks. **3)** Media is transferred into glass vials and diluted chemical is added. **4)** T-25 filter caps are replaced with air-tight caps with septums and cell media containing chemicals is added via syringe and left for 24 hours. **5)** After 24 hours, chemical-containing media is removed and cells are washed with buffer. **6)** T-25 air-tight septum caps are replaced with filter caps and cells are cultured for an additional 24-hour “chase” period.
