## Supplementary file 26 for "Endocrine disruptor-induced epimutagenesis *in vitro*: Insight into molecular mechanisms"

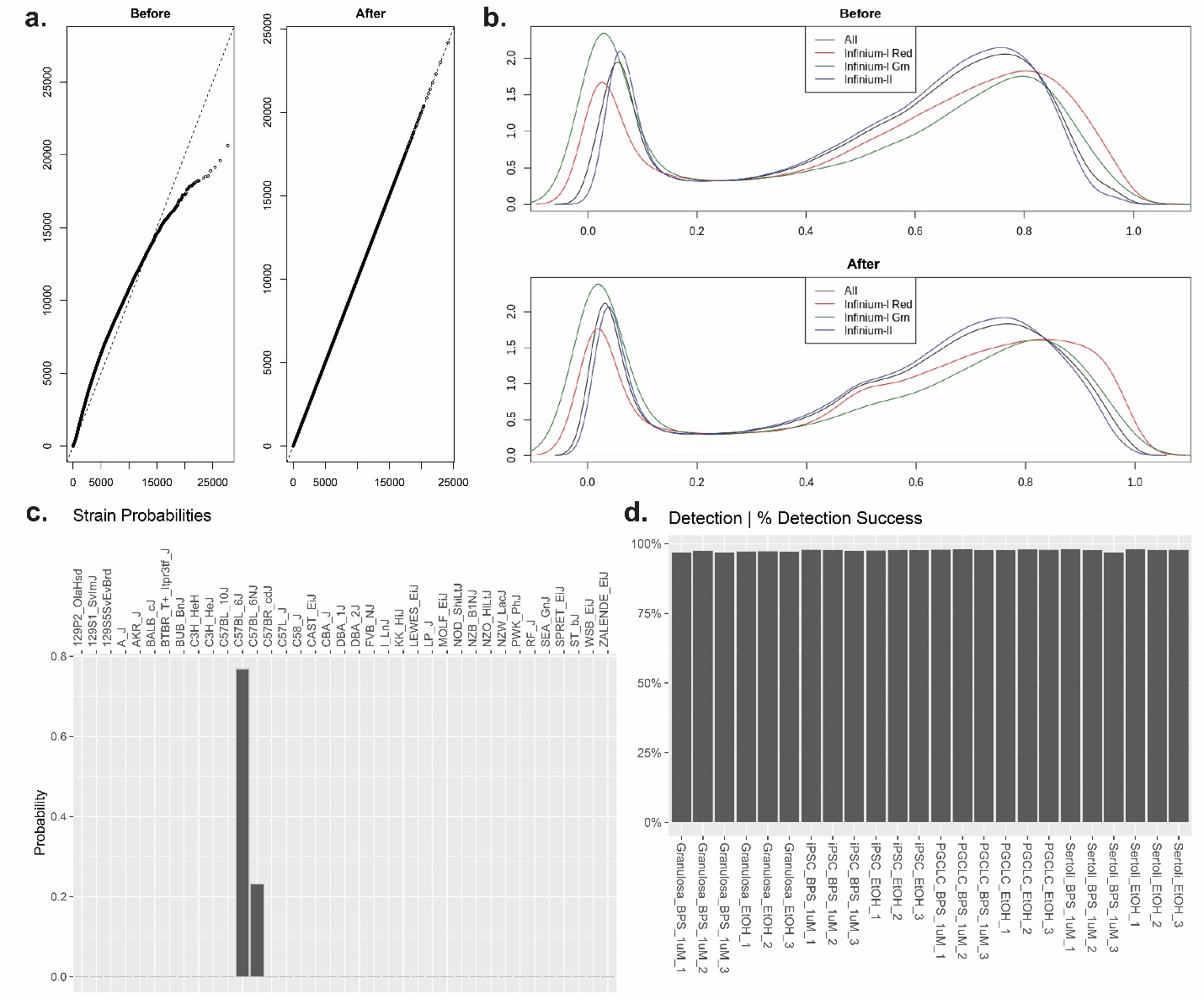

**Quality control for Infinium Mouse Methylation BeadChip Array data. a)** Non-linear correction of dye bias removal from sample data. **b)** Background signal subtraction from samples to limit noise. **c)** Prediction of correct C57B6 mouse strain from samples included in the study based on built-in controls on Infinium Mouse Methylation BeadChip Array. **d)** Average CpG probe detection success of 97.58% percent across all samples indicating efficient bisulfite conversion of all samples.
