## Supplementary file 27 for "Endocrine disruptor-induced epimutagenesis *in vitro*: Insight into molecular mechanisms"

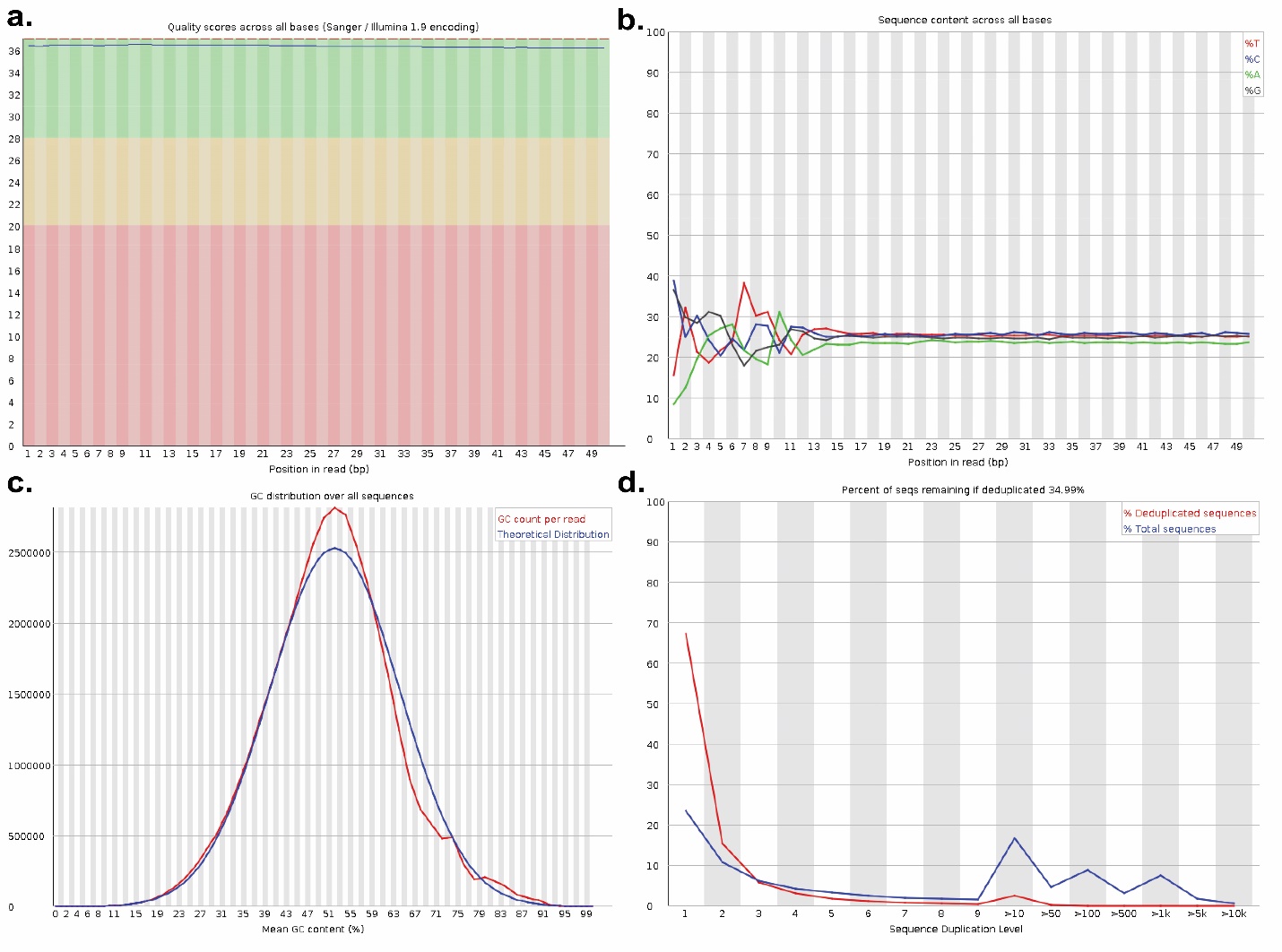

**Quality control for RNA-seq data.** RNA-seq quality control data from one of the three PGCLC replicates exposed to BPS as an example. **a)** Base calls showed high quality scores (phred scores > 30) for all bases in reads. **b)** Reads showed equal distributions of all four bases following the initial adaptor sequence and sufficient base complexity. **c)** Distribution of GC sequences across reads aligned very closely with the theoretical distribution. **d)** Duplication plot indicates deduplicated libraries contained ~67% unique sequences which indicates sufficient library complexity for subsequent downstream data processing.
