## Supplementary file 28 for "Endocrine disruptor-induced epimutagenesis *in vitro*: Insight into molecular mechanisms"

**Quality control metrics for EM-seq data.**

| Sample | Insert Size (bp) | Median Coverage | Read Counts (Millions) | Aligned  Reads  (%) | Duplicate Reads  (%) | GC  (%) |
| --- | --- | --- | --- | --- | --- | --- |
| iPSC_E_1 | 320 | 37 X | 851.9 | 99.0 | 28.6 | 22 |
| iPSC_E_2 | 374 | 33 X | 752.6 | 99.1 | 26.4 | 22 |
| iPSC_E_3 | 378 | 38 X | 873.5 | 99.2 | 27.2 | 22 |
| iPSC_BPS_1 | 360 | 28 X | 626.7 | 99.1 | 23.6 | 22 |
| iPSC_BPS_2 | 330 | 30 X | 695.3 | 99.1 | 26.6 | 22 |
| iPSC_BPS_3 | 338 | 33 X | 755.8 | 99.1 | 27.7 | 22 |
| PGCLC_E_1 | 388 | 32 X | 707.9 | 99.0 | 25.6 | 23 |
| PGCLC_E_2 | 374 | 25 X | 559.4 | 99.0 | 24.0 | 22 |
| PGCLC_E_3 | 412 | 30 X | 673.5 | 99.2 | 24.9 | 22 |
| PGCLC_BPS_1 | 399 | 31 X | 672.9 | 99.1 | 25.6 | 22 |
| PGCLC_BPS_2 | 371 | 32 X | 753.3 | 99.1 | 28.6 | 22 |
| PGCLC_BPS_3 | 392 | 27 X | 599.3 | 99.1 | 23.6 | 22 |
